# sCNAphase: using haplotype resolved read depth to genotype somatic copy number alterations from low cellularity aneuploid tumors

**DOI:** 10.1101/038828

**Authors:** Wenhan Chen, Alan J. Robertson, Devika Ganesamoorthy, Lachlan J.M. Coin

**Affiliations:** Institute for Molecular Bioscience, The University of Queensland, St Lucia, Queensland, 4072, Australia

## Abstract

Accurate identification of copy number alterations is an essential step in understanding the events driving tumor progression. While a variety of algorithms have been developed to use high-throughput sequencing data to profile copy number changes, no tool is able to reliably characterize ploidy and genotype absolute copy number from tumor samples which contain less than 40% tumor cells. To increase our power to resolve the copy number profile from low-cellularity tumor samples, we developed a novel approach which pre-phases heterozygote germline SNPs in order to replace the commonly used ‘B-allele frequency’ with a more powerful ‘parental-haplotype frequency’. We apply our tool - sCNAphase - to characterize the copy number and loss-of-heterozygosity profiles of four publicly available breast cancer cell-lines. Comparisons to previous spectral karyotyping and microarray studies revealed that sCNAphase reliably identified overall ploidy as well as the individual copy number mutations from each cell-line. Analysis of artificial cell-line mixtures demonstrated the capacity of this method to determine the level of tumor cellularity, consistently identify sCNAs and characterize ploidy in samples with as little as 10% tumor cells. This novel methodology has the potential to bring sCNA profiling to low-cellularity tumors, a form of cancer unable to be accurately studied by current methods.

## INTRODUCTION

Somatic Copy Number Alterations (sCNAs) represent an important class of mutation in the cancer genome, evident by the large number of short focal sCNAs and larger chromosomal scale changes seen in the analysis of individual tumor genomes (1). This class of mutation has been linked to tumor progression, metastasis, multidrug resistance and poor clinical outcomes (2-6). Despite the sporadic accumulation of sCNAs during tumor progression, a number of regions are subject to recurrent sCNAs (7). Some of these recurrent sCNAs are found across different cancer types, while others were specific to a particular type or subtype of the disease (6, 8-10). As a result, determining the sCNAs in an individual tumor sample has become standard practice in pathology labs for the treatment of some cancers. For example, this type of analysis is routinely used to assign the optimal chemotherapeutic treatments for patients with breast cancer who contain additional copies of the *HER2* gene (11, 12).

Despite the importance of this class of mutation, it can be difficult to characterize the copy number profile of a tumor genome (13). A typical tumor biopsy will contain both tumor cells as well as cells with a normal, diploid genome. This can be quantified via the cellularity (the proportion of tumor cells in this mixture) or via the tumor DNA purity (the proportion of tumour DNA in the mixture of normal and tumor DNA). Tumour purity is a function of both the cellularity and the tumor ploidy (which we define as the average copy number of the tumor) - for example a 50% cellularity tetraploid tumour and will have a 66% tumor purity. The best current methods fail to produce the correct copy number segmentation when tumor cellularity in a sample falls below 40% (**Supplementary Table S1**). The cellularity for a number of serious forms of cancer, such as Breast Invasive Carcinoma, Lung Adenocarcinoma and some forms of Melanoma routinely fall below this threshold (13), moreover multiple cancers including renal clear cell carcinoma and lower grade glioma show a decreased survival time with lower tumor cellularity (14). Thus there is an important unmet need to identify copy number mutations in low purity samples.

It is possible to survey the copy number profile of samples with a high tumor content, using a number of different techniques, however it is difficult to characterize the full spectrum of sCNAs with a single technology. Spectral Karyotyping (SKY) is one of the most accurate tools for characterizing and visualizing genome wide changes in ploidy (15-18), but suffers from a limited resolution and is low throughput. More recent technologies, such as single-nucleotide polymorphism (SNP) microarrays has provided powerful approaches for interrogating the tumor genome and identifying copy number mutations (13, 19-22). These include ASCAT (23) and ABSOLUTE (13), both of which initially pre-segment SNPs into regions of equal copy-number (using a threshold based, model-free approach) and subsequently estimate ploidy and tumour purity by use of a model for the observed read-depth data conditional on the fixed segmentation. ASCAT and ABSOLUTE are highly successful in samples with as little as 40% tumor DNA (13, 23), however the reliance on an initial model-free segmentation is likely to limit the ability of these methods to detect copy number alterations at lower tumour cellularities (**Supplementary Table S1**). The performance of these tools are also restricted by the resolution of different microarray platforms as well as fluorescence signal saturation at high copy number.

High throughput sequencing (HTS) is a powerful technology for identifying sCNAs, which may make it possible to characterize the complete copy number profile of impure cancer samples. Both ASCAT and ABSOLUTE have been modified to be applicable to HTS, and a number of other computational tools have been specifically designed to identify copy number changes, characterize loss of heterozygosity (LOH) and identify homozygous deletions from HTS data (**Supplementary Table S1**). These tools use a variety of different signals present in HTS data including read depth aberration, B-allele frequency at somatic and germline SNPs. Most of these tools are not suitable for samples with tumor cellularity less than 40% (23-26). CLImAT is a recently introduced tool, which uses read-depth and BAF to estimate the ploidy and purity of impure tumor genomes and also characterizes copy number and LOH changes (27). At 20% simulated tumor cellularity, CLImAT demonstrated more robust cellularity and ploidy estimates as well as greater sCNA and LOH calling accuracy than Absolute (13), SNVMix (24), Control-FREEC (25) and Patchwork (26); however this was evaluated using simulated tumor chromosomes rather than tumor-normal mixture samples.

Modeling BAF has strengths and limitations complementary to read-depth (RD) modeling. BAF - which effectively uses an internal control of one allele versus the other - is less susceptible to position-specific biases, such as GC and mappability biases. However, one striking disadvantage of BAF modeling is that it is not possible to directly summate the allelic depth signal over multiple adjacent SNPs, as the non-reference alleles at one adjacent position may not be on the same parental haplotype. Summating RD over windows of size from 10kb up to 1Mb leads to substantial increases in statistical power to detect sCNAs. We hypothesized that application of state-of-the-art computational phasing approaches, incorporating population haplotype from the 1000 genomes project (27) as well as direct within-read phasing (28), would allow us to sum allelic depth along phased haplotypes to obtain *parental-haplotype frequency* (PHF) estimates. We further hypothesized that modeling PHF instead of BAF could lead to improved power characterize tumor ploidy and sCNAs at ultra-low levels of tumor purity.

In this manuscript we present sCNAphase (https://github.com/Yves-CHEN/sCNAphase), a tool, which has been developed to characterize the full copy number profile of a cancer sample across a range of tumor purity. It achieves this by inferring tumor ploidy, sCNAs and regions of LOH across all levels tumor purity by integrated modeling of PHF and RD. We show that sCNAphase has accuracy comparable to SKY in determining genome-wide changes in ploidy and is able to identify focal sCNAs that are consistent with results from microarray analyzes. We also show that sCNAphase can confidently determine regions that have undergone a loss of heterozygosity event and identify regions of homozygous deletion. Moreover, sCNAphase consistently generates accurate sCNA segmentations at low levels of tumor purity and can accurately define levels of tumor purity in mixtures containing 5% tumor DNA.

## MATERIAL AND METHODS

### Datasets

The Illumina whole genome sequencing data of four pairs of tumor and matched normal cell line samples were downloaded from Cancer Genome Atlas (TCGA) (29) or Illumina BaseSpace (https://basespace.illumina.com). All of the 8 samples are at higher than 49x coverage, expect for HCC2218BL at 37x (Table 1). Two independent normal 30X samples were also available for HCC1143 and HCC1954. For the other two samples HCC1187 and HCC2218, we generated a second ‘normal’ by downsampling the available matched normal to 30X. We will refer to these second normal samples as ‘0% mixtures’ as we will use them as a negative control to investigate whether methods detect tumor DNA in mixtures without tumour DNA.

**Table 1.** Information about the cell-line samples.

| Cell-line | Platform/<br>Library | Downloaded from | Coverage | Tissue | Annotation |
| --- | --- | --- | --- | --- | --- |
| **HCC1143** | HiSeq2000/<br>Paired WGS | TCGA | 50x | Breast<br>ductal | 52 years female, Caucasian with STAGE IIA,<br>grade 3 Breast ductal carcinoma<br>Basal A subtype |
| **HCC1143BL** | HiSeq2000/<br>Paired WGS | TCGA | 60x, 30x | Blood | Paired 60x normal for HCC1143 and<br>independent 30x sample |
| **HCC1954** | HiSeq2000/<br>Paired WGS | TCGA | 58x | Breast<br>ductal | 61 years female, Indian with STAGE IIA, grade<br>3 Breast ductal carcinoma.<br>Basal A subtype, HER2 amplified |
| **HCC1954BL** | HiSeq2000/<br>Paired WGS | TCGA | 71x, 30x | Blood | Paired 60x normal for HCC1954 and<br>independent 30x sample |
| **HCC1187** | HiSeq2000/<br>Paired WGS | Illunima BaseSpace | 93x | Breast<br>ductal | 41 years female, Caucasian with STAGE IIA,<br>grade 3 Breast ductal carcinoma<br>Luminal |
| **HCC1187BL** | HiSeq2000/<br>Paired WGS | Illunima BaseSpace | 49x | Blood | Paired normal for HCC1187 |
| **HCC2218** | HiSeq2000/<br>Paired WGS | Illunima BaseSpace | 83x | Breast<br>ductal | 38 years female, Caucasian with STAGE IIIA,<br>grade 3 Breast ductal carcinoma<br>Basal A subtype, HER2 amplified |
| **HCC2218BL** | HiSeq2000/<br>Paired WGS | Illunima BaseSpace | 37x | Blood | Paired normal for HCC2218 |

By mixing the different amount of reads from pure tumor samples together with the ‘0% mixture’ samples (i.e. the second normal sample), a series of mixtures samples were created at 30x coverage with 5%, 20%, 40%, 60%, 80% and 95% of tumor DNA. These mixture samples were obtained as BAM files from the Cancer Genome Atlas (TCGA) (29) for HCC1143 and HCC1954, from Illumina BaseSpace for HCC1187 and HCC2218 (**Supplementary Table S2**). We also created an extra 10% mixture for all samples.

### Phasing Matched Normal

By running samtools (30) and BCFtools (30) on the normal samples, we determined the germline heterozygous SNPs. At these loci, samtools is used to calculate the depths for the tumor sample. The somatic mutations were ignored in this step. Then SHAPEIT2(28), an *in silico* haplotype phasing tool, was used to phase whole genome sequence short read data. Because SHAPEIT2 requires a set of pre-phased reference haplotypes as input, we used pre-phased haplotypes provided by SHAPEIT2 calculated from the 1000G Phase I dataset. Based on this analysis, we assigned each allele from each heterozygote in the matched normal to a haplotype, labelled H1 or H2.

### Calculating regional haplotype depth

Our approach only considers read-depth at heterozygous germline SNPs, and ignores information from somatic mutations. We calculate the total read depth (RD) for the tumour (t) and normal (n) in windows i = 1..N, each consisting of *K* germline heterozygous SNPs (with a default *K =* 40) as

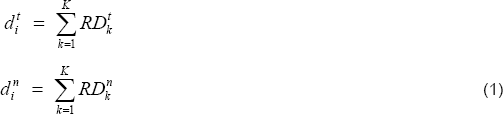

We also calculate the read-depth in these regions specific to haplotype H1 (RD_H1_) as

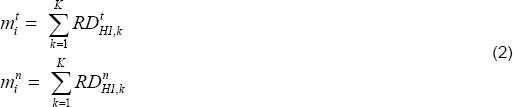

If the *K* SNPs in a window are split by a large gap (greater than 1M, e.g a centromere), this window is excluded from further analysis. We discuss below detection of copy number switches which occur within a window of n SNPs.

The data modelled by sCNAphase is **D** = (**m**^t^**, d**^t^**, m**^n^**, d**^n^), where **m**^t^ = {m^t^_i_} **d**^t^ = {d^t^_i_}, **m**^n^ = {m^n^_i_} and **d**^n^= {d^n^_i_}. We also calculate the values D^t^, D^n^ as the sum totals of **d**^t^**, d**^n^ respectively across the genome.

We also define the tumour *parental-haplotype frequency* (PHF^t^), which is a generalisation of the standard B-allele frequency (BAF) as

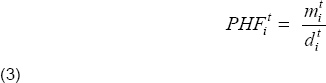

### Statistical model of haplotype depth under null hypothesis of absence of tumor DNA

Under the null hypothesis of the presence of no tumor DNA in the sample or the matched normal the distribution of H1 allele counts *m*_i_^t^ calculated at each window *i* can be modelled with a binomial distribution with a 50% ‘probability of success’

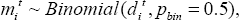

However, the presence of copy number variation and mapping biases lead to shifting from 0.5 to an unknown *p*_0_ and greater than expected variation in *m*_i_^t^. As a result, it is necessary to instead use a beta-binomial with parameters alpha and beta equal to the number of counts mapping to H1 and H2 in the normal sample:

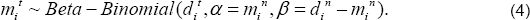

As *m*_i_^n^ and *d*_i_^n^ − *m*_i_^n^ increase, the beta-binomial distribution approaches to the Binomial(*m*_i_^t^, *d*_i_^t^, p_bin_=p_0_).

### Tumor purity and cellularity

The percentage of tumor content can be measured in two different scales, 1) tumor cellularity (tc) defined as the percentage of tumor cells or 2) tumor purity (*tp*) defined as the percentages of tumor DNA in mixtures of tumor and normal cells. These two quantities are related via the tumor ploidy, which we define as the average copy number of the tumor over all windows:

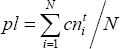

If we assume a diploid normal, then the relationship between tumor cellularity (tc) and tumor purity (tp) is given by:

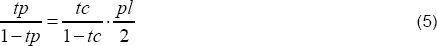

### Hidden Markov Model based on haplotype segments

We model the probability of the data conditional on the tumor celullarity *tc* and the tumor ploidy *pl* as

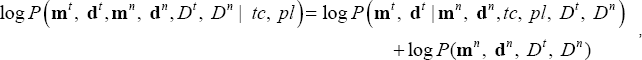

assuming that the normal depth is independent of tumor cellularity and ploidy, and the total tumor read depth is also independent of cellularity and ploidy.

We use a hidden Markov Model to calculate this probability. The hidden states (*s*_i_) of this model are the unobserved copy numbers, x for H1 and y for H2 in the tumor genome in window i, represented by g = (x, y). The total copy number is given by CN(*g*) = x + y. We consider all hidden states *g* with copy number in the range 0<=*CN*(*g*)<=12, that makes up a set ***G*** of 91 possible configurations. We also define a transition probability for transitioning between pairs of states *g* and *l* as t(s_i_ = g | s_i-1_ = *l*). We can write down the joint probability of the observed tumor depth data and the unobserved state path using the following equation:

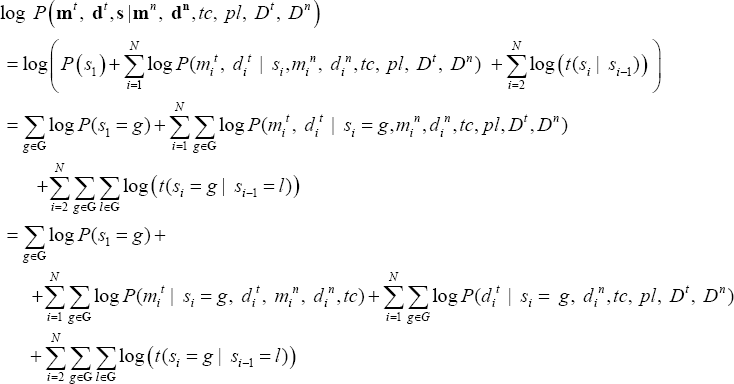

#### Emission probability

In the above equation, the emission probability was split into two parts (the second and third components): one models the effect of sCNA on the total read depth *d*_i_^t^ comparing to D^t^ and another models the effect of sCNA on read depths for H1 comparing to H2. They can be calculated as,

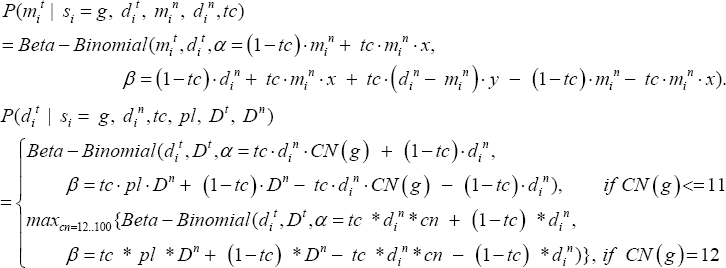

This effectively extends the maximal copy number to 100 to capture the significantly amplified regions, e.g. 50 copies of *HER2*, which otherwise seems equally unlikely to be any of the genotypes with copy number <= 12.

#### Initial and transition probability

The initial probability p(s_1_= *g)* in **Equation 6** was set to be 1/91 for each *g* from **G**, so that show no preferences for any genotypes. We used a fixed transition probability model defined on the all the states G at locus *i*. The transition probability two successive sites s_i-1_ and s_i_ is defined as

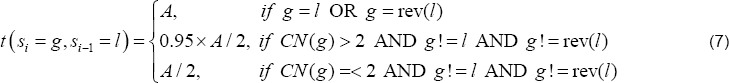

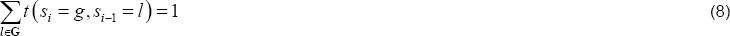

in which, rev(*l*) is to reverse the order of the copy numbers for H1 and H2 for state *l*. Combining Equation (7, 8), *A* can be calculated. Essentially, this transition probability imposes a weak preference for lower copy number states.

#### Estimation of tumor cellularity and ploidy

We first estimate the tumor purity and ploidy by maximizing the likelihood function in **Equation 6** calculated by Baum-welch algorithm (31). Because of the flat likelihood surface at low purity tumor (**Supplementary Figure S1**), two dimensional gradient-free local searching algorithms occasionally failed to find the global maximum. We solved this by performing a one dimensional optimization to determine the ideal tumor cellularity, for a set of ploidy values increased from 1.8 to 5 with an increment of 0.2. The pair of cellularity and ploidy with the biggest likelihood are chosen as cellularity and ploidy estimate.

#### Estimation of copy number profile

With the estimated tumor purity and ploidy, the hidden states (***s***) at each the window *i* are estimated using Viterbi algorithm(32). The density of the windows calculated by counting the distance between two neighbor windows (**Supplementary Figure 2**) is less than 100kb for the vast majority. To impute the copy number information for regions in between two successive windows (segments), two windows are joined into one segment, if they are with the same copy number state and locate in short distance of less than 100kb. This generates a copy number segmentation for the whole genome, with a few undetermined regions when two windows cannot be joined.

### Removal of merging errors at copy number switches

The merging of every *K* SNPs gives a merging error, if the region spans the boundary of two adjacent sCNAs, which cannot be resolved with a single copy number state. Therefore we nullify the estimation for segments with merging errors, which can be detected by a likelihood ratio test for each SNP *k* in a window *i* as follows,

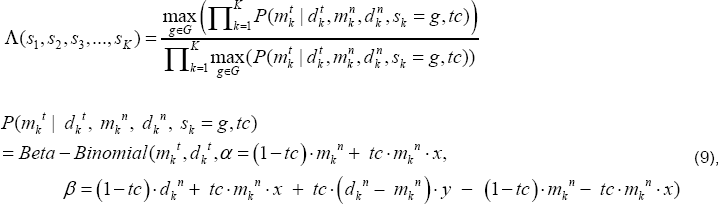

in which m_k_^t^, d_k_^t^ are the alternative allelic depth and the read depth from a tumor sample at locus k; m_k_^n^ and d_k_^n^ are depths from the matched normal sample. The numerator assumes the *K* SNPs are in a single sCNAs, with a single copy number state *g* from *G* which maximizes the likelihood. The numerator assumes each SNP *k* can have an individual genotype that maximizes the likelihood. We then removed 1% of the segments with minimal values.

### Workflow

We built up a pipeline – sCNAphase to generate a full copy number profile including copy number alterations to each parental haplotype, tumor purity and tumor ploidy using the method described above.

**The pre-processing steps** of the workflow (**Figure 1A**) include 1) calling germline SNP variants, 2) resolving germline allelic haplotypes and 3) generating regional haplotype depths. Compared with B-allelic depth at per site, the phased allelic depths generated by ordering the alleles according to parental haplotypes (H1 and H2), gives a less variable allelic frequency, based on which the regional haplotype depths further reduce the variability (**Figure 1B, C, D**). The regional haplotype depth information is then used for the estimation step.

**Figure 1.**
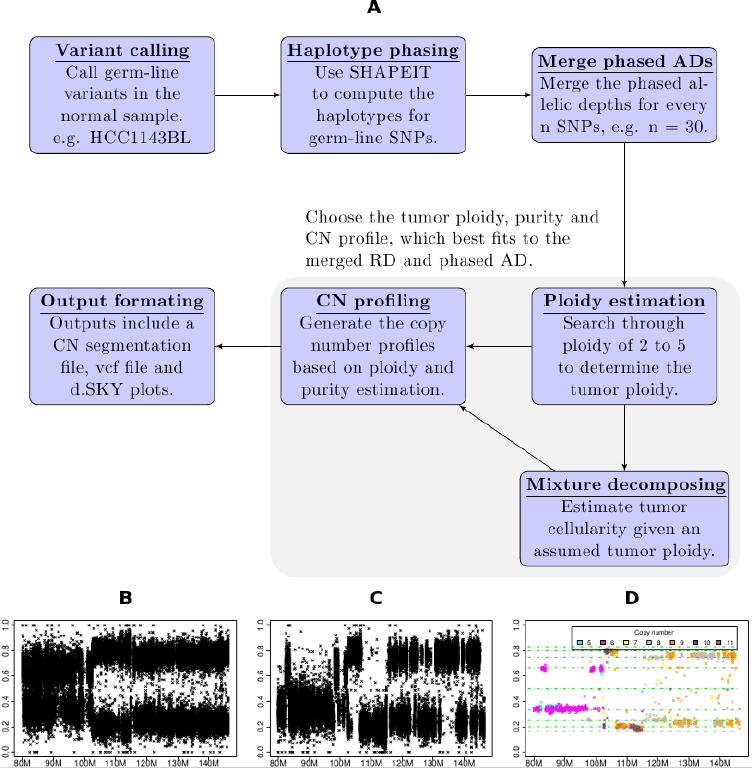
The individual steps that make up the sCNAphase workflow. (A) Each panels shows an individual process used by sCNAphase to characterize the copy number profile from matched tumor and normal pairs. (B) The BAFs from a region of chromosome 8 from the HCC1143 breast cancer cell-line. (C) The application of phasing to the BAF data makes it possible to identify PHFs, regions composed of 40 adjacent germline, heterozygous SNPs. Each PHF increase the power of this analysis and makes it possible to better reflect the copy number profile of this region. (D) Application of the sCNAphase pipeline to the phased data, calls specific copy number changes.

**The estimation step** performed an integrated calculation of tumor purity, tumor ploidy and copy number profile, using the idea from **Equation 6**. The output from this includes a) a copy number segmentation file that shows the regional changes in overall copy number as well as haplotype copy number in a region; b) a digital SKY (dSKY) plot based on the segmentation is generated for visualization similar to traditional SKY plot; c) a vcf file includes allelic copy numbers and phases for the each germline SNPs. This estimation procedure was implemented in R, powered by multithreading techniques. Excluding the pre-processing, the estimation takes approximately 4 hours running on 12 CPUs (1600MHz) for the Illumina whole genome sequencing data.

### Comparison of two copy number segmentations

Using one segmentation as the reference, the consistency of the other segmentation (test segmentation) with the reference can be measured in varied ways, given different criteria for counting if two copy number states are consistent. We performed this analysis using three different criteria:

#### Counting per base overlap for copy number gain or loss

A segment is identified as gain if the copy number is higher than the average ploidy; otherwise, it is identified as loss according to a particular segmentation. The two segmentations are counted as overlapped at a single base, if that base is consistently seen as gain or loss. The fraction of overlapped bases to the number of bases in reference and the test segmentation are defined as sensitivity and specificity respectively

#### Counting per base overlap for LOHs

The sensitivity and specificity are calculated based on how many bases are consistently identified as loss of heterozygosity. This only stresses identification of one haplotype copy number being zero (not both), disregards whether the overall copy numbers from two predictions being equal.

#### Counting 50% reciprocal overlap for focal amplifications

A segment is identified as focal amplification if the copy number is at least twice of the average ploidy and the size is between 100kb to 4Mb. Once a segment is identified as a focal amplification, the actual copy number is disregarded. A reference segment and a test segment are overlapped, if the overlapped region accounts for at least 50% of each of the two segments. The total number of overlapped segments to the number of segments from reference or test segmentation are defined as the sensitivity and specificity respectively.

## RESULTS

We developed a new method, sCNAphase, for estimating the copy number and LOH profile of the tumor genome for low cellularity tumors. sCNAphase integrates haplotype-specific allele counts together with total read depth in a Hidden Markov model which explicitly models both tumor DNA purity and ploidy. We evaluated the performance of sCNAphase against two state-of-the-art algorithms (CLImAT (version 1.1) and ASCAT (version 2.4)) using mixture samples derived from whole genome sequence data from 4 well-characterized tumor cell-line samples covering a range of ploidies with known copy number information from SNP array and SKY data. The samples used in this study are described in Table 1. Mixture samples were generated *in-silico* over a range of tumor purities from 5% to 95% as described in Methods.

### Haplotype phasing greatly improves the power to identify sCNA from allelic depth

The B-allele frequency (BAF) is commonly used to detect the presence of copy number variants (CNVs) in normal, diploid genome(33). This signal can also be interrogated to find sCNAs by looking for deviations from an expected equal ratio of two alleles at germ-line heterozygous SNPs. In order to investigate the power of this signal, we calculated Allelic Depth (AD) at all germ-line heterozygous SNPs in a tumor derived cell-line (HCC1143, see Table 1) at varying tumor purities, as well as in the matched normal sample (note that an independent normal was used to generate artificial mixed tumor/normal samples at varying purity, see Methods). The distribution of p-values under a null model which assumes an equal ratio is inflated even for the normal sample, due to the presence of germline CNVs (**Supplementary Figure S3**), thus we instead calculate p-values under the assumption that tumor BAF is the same as germ-line BAF, which corrects this inflation (**Figure** 2A). BAF provides substantial power to identify sCNA at 100% purity (**Figure** 2B.), but the signal is too weak at 5% purity to identify any sCNAs (**Figure** 2C.).

**Figure 2.**
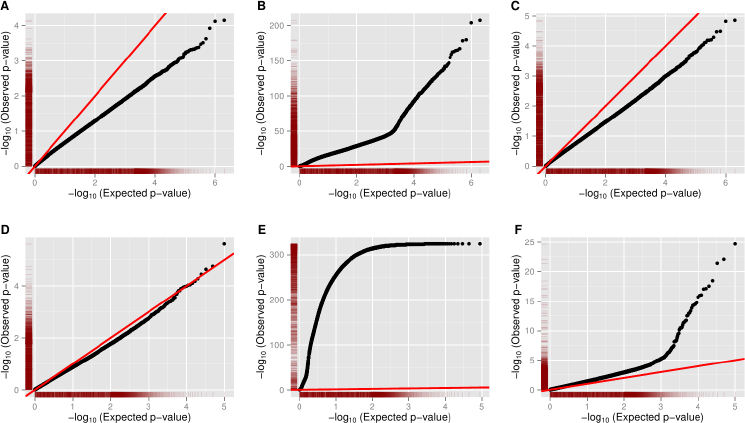
The capacity of different allele frequencies to determine sCNAs in tumor derived cell-line samples. Each of the points in the Q-Q plots corresponds to a P-value for a pair of allelic depths from a tumor derived cell line (HCC1143) at all germ-line heterozygote SNPs in different scenarios. The red line represents the expected distribution of p-values under the null hypothesis (of no sCNA). Pvalues are calculated using a binomial distribution with probability of success equal to observed germline BAF (A, B, C) or PHF (D, E, F). Deviation above this line indicates power to detect sCNA (A) Using BAF on 0% tumor purity sample (i.e. normal). (B) Using BAF at 100% tumor purity (C) Using BAF at 5% tumor purity. (D) Using PHF on a 0% tumor purity sample (i.e. normal). (E) Using PHF at 100% tumor purity. (F) Using PHF on 5% tumor purity.

To investigate the improvement in power to detect sCNAs using PHF rather than BAF, we phased germ-line heterozygotes using SHAPEIT2 (28) and the panel of reference haplotypes from the 1000 Genomes Project, then used this information to calculate tumor PHF in non-overlapping windows of 40 consecutive heterozygous SNPs (Methods). This strategy dramatically improved the power to detect sCNAs in the 100% purity sample (**Figure** 2E) and 5% (**Figure** 2F), and did not lead to spurious identification of sCNAs in the normal sample (**Figure** 2D).

### Digital Spectral Karyotyping with sCNAphase is concordant with spectral karyotyping

The genome-wide results from the analysis of each of the pure tumor cell-lines and the matched germline samples were visualized using *digital spectral karyotyping* (dSKY) plots. These images were designed to build on the effectiveness of spectral karyotyping (SKY) images to represent genome wide changes in ploidy (**Figure** 3). Our dSKY plots make visual identification of aneuploidy, loss of heterozygosity and focal changes straightforward. To build on SKY platform, red shading on the chromosome ideograms was added to indicate regions of LOH, while green shading indicates homozygous deletions.

**Figure 3.**
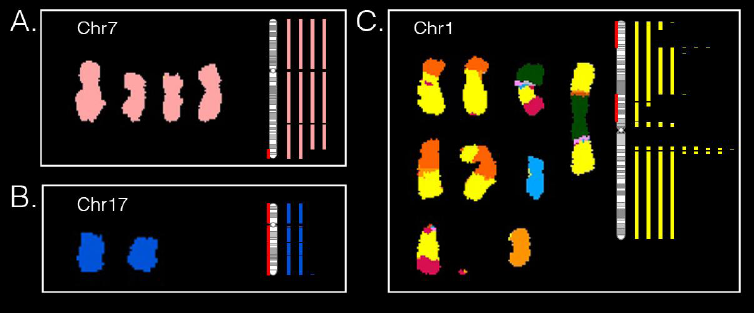
dSKY plots build on SKY images to better reflect the complete copy number profile of a tumor genome. In each of these subplots (A, B, C), a SKY image (reproduced with permission from (52)) on the left shows the copy number of a particular chromosome, which was compared to a dSKY plot of the same chromosome - the vertical bars shown on the right in the same color. The number of the bars correspond the copy numbers at the specific regions calibrated by the chromosomal ideogram (vertical bar with grey bandings). The red shading on the ideogram is used to mark regions of LOHs.

Comparing the 3 available SKY images from these cell-lines to the corresponding dSKY plots, demonstrated that the results from sCNAphase were highly concordant with those from the SKY analyses, both at the ploidy level (**Table** 2) and at the large-scale genomic alterations level. For example, both the sCNAphase and SKY analysis of HCC1187 classified this cell-line as hypo-triploid and both suggested that this cell-line had 4 copies of chromosome 7, and 2 copies of chromosome 17 (**Figure 3AB**; **Supplementary Figure S4**). In addition to the chromosomal gains or loss shown in SKY, dSKY plots are able to display more important information such as focal copy number changes, regions with loss of parental chromosomes and regions that are altered by events more complicated than whole chromosome gains or loss. For example, at the chromosome level, both SKY and sCNAphase suggest that HCC1187 has 4 copies of chromosome 7 (**Figure 3A**). However, the higher resolution result from sCNAphase was able to detect that two of these copies shared an identical deletion at the tip of the q arm. Analysis of chromosome 17, revealed that while both the SKY and the dSKY suggested HCC1187 contained two copies of the chromosome (suggesting this chromosome had not undergone a copy number mutation), the dSKY plot revealed this chromosome has undergone a copy neutral loss of heterozygosity (LOH) event (**Figure 3B**).

**Table 2.** Summary of results from sCNAphase on 4 cancer cell lines. The proportion of the genome that has undergone sCNA is calculated as the proportion of the genome with copy number not equal to nearest integer ploidy.

| Cell-line | Estimated Ploidy |  | Proportion of genome |  |  |
| --- | --- | --- | --- | --- | --- |
|  | Ploidy determined by sCNaphase | Ploidy determined by COSMIC | Undergone sCNA | Undergone LOH | Undergone LOH with more than 2 copies |
| **HCC1143\***<br>Hypo-tetraploid | 3.7 | 3.36 | 83% | 46% | 30% |
| **HCC1954\***<br>Tetraploid | 4.5 | 4.2 | 43% | 8% | 6% |
| **HCC1187\***<br>Hypo-triploid | 2.7 | 2.64 | 28% | 58% | 8% |
| **HCC2218#**<br>Tetraploid | 4.2 | 3.93 | 64% | 14% | 4% |

\* Cell-line with the ploidy determined from SKY (52).

In addition to visualizing large-scale chromosomal copy number alterations, inter-chromosomal translocations make it difficult for traditional SKY analyses to determine the exact copy number of translocated regions. For example, the results of from the SKY analysis of chr1 in HCC1187, make it difficult to deconvolute the copy number profile of this chromosome at the genome level (**Figure 3C**). While the dSKY plots do not provide any information about the translocations, they are able to demonstrate which region has undergone a specific copy number alteration. Together these examples show that dSKY plots, make it possible to begin to untangle complex phenotypes and identify mutations invisible to previous methodologies at a genome-wide level.

### sCNAphase accurately calculates tumor ploidy, tumor purity and sCNAs across a range of different levels of simulated tumor purity

The presence of stromal cells or other cells with a normal diploid genome in a solid tumor sample can impact the capacity of genomic-based approaches to characterize mutations in a tumor sample (13). To assess the performance of sCNAphase to characterize impure tumor samples, simulated mixtures from HCC1143 and HCC1954 (29) as well as HCC1187 and HCC2218 from Illumina BaseSpace (https://basespace.illumina.com) as well as HCC1187 and HCC2218 from Illumina BaseSpace (https://basespace.illumina.com) were analyzed (see Methods). To simulate the range of tumor heterogeneities found in primary tumor samples, each cell-line had a number of mixtures analyzed, each with varying proportions of tumor cell-line DNA (95%, 80%, 60%, 40%, 20%, 10%, 5% and 0% - see methods). Each mixture and matched normal sample were passed through the sCNAphase pipeline. Given that the ploidy estimates from the pure cell lines were comparable to the results from SKY (**Table** 3), we assessed the degree to which the analysis of the mixture samples were consistent with the results from the pure cell lines. A similar analysis was carried out using CLImAT and ASCAT, which estimate sCNAs as well as tumor purity and ploidy.

**Table 3.** The capacity of sCNAphase to detect the focal sCNAs in COSMIC

| Tumor purity | HCC1143<br>15 focal amplification from COSMIC |  | HCC1954<br>94 focal amplification from COSMIC |  | HCC1187<br>2 focal amplification from COSMIC |  | HCC2218<br>18 focal amplification from COSMIC |  |
| --- | --- | --- | --- | --- | --- | --- | --- | --- |
|  | Sen | Spe | Sen | Spe | Sen | Spe | Sen | Spe |
| **100%** | 80 | 23 | 66 | 52 | 100 | 26 | 61 | 31 |
| **80%** | 80 | 24 | 67 | 47 | 100 | 25 | 56 | 31 |
| **60%** | 80 | 24 | 63 | 47 | 100 | 27 | 44 | 38 |
| **40%** | 67 | 28 | 61 | 52 | 100 | 30 | 78 | 25 |
| **20%** | 80 | 22 | 65 | 57 | 100 | 30 | 61 | 44 |
| **10%** | 87 | 29 | 77 | 52 | 100 | 23 | 61 | 48 |
| **5%** | 80 | 23 | 37 | 93 | 100 | 29 | 22 | 100 |

Sen for sensitivity; Spe for specificity.

This analysis revealed that sCNAphase was able to accurately recapitulate the ploidy results from each of the pure cell-lines across the majority of the mixtures as well as accurately determine the level of tumor DNA in each sample (**Figure** 4AB, **Supplementary Table** S3). The only inaccurate ploidy calls came from HCC1954 and HCC2218 mixtures at 5% tumor purity. Despite this, sCNAphase was still able to accurately report on the amount of cell-line DNA in these samples. Across the entire cohort of mixtures, the sCNAphase results only deviated from the simulated proportion by maximum 2% (**Figure** 4B, **Supplementary Table** S3). Cellularity estimates, which can be calculated as a function of tumor purity and ploidy (see Methods) were also reported (**Figure** 4C). The sCNAs identified were consistent across the mixtures (see Methods) except for the 5% mixtures from HCC2218 and HCC1954 in which the ploidy estimates had been incorrectly calculated (**Figure** 4D). Adjusting the ploidy to the correct value rescued the segmentation and allowed sCNAphase to capture the same broad copy number profile from these 5% mixtures (**Supplementary Figure** S5).

**Figure 4.**
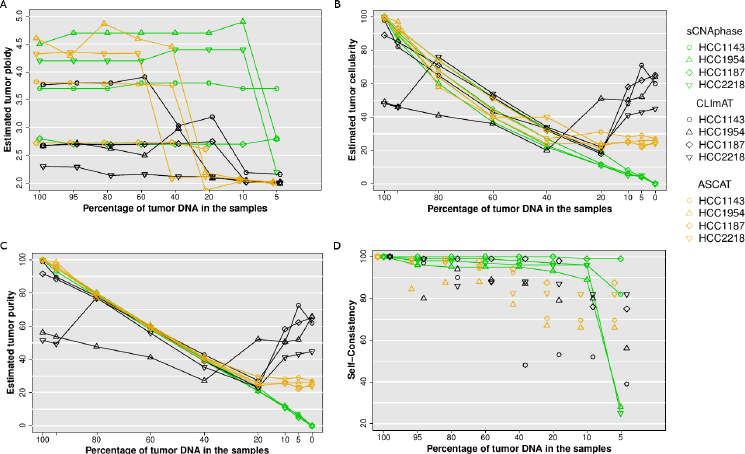
(A) Inferred ploidy as a function of tumor purity. (B) Estimated tumor cellularity (the proportion of tumor cells in each sample) at different levels of simulated purity. (C) Estimated tumor purity versus simulated tumor purity (proportion of tumor DNA in sample). These purity values were calculated from estimated cellularity and estimated tumour ploidy using Equation 6 for each tool. (D) Self-consistency at decreasing tumor purity, measured as average of base-pair sensitivity and basepair specificity versus 100% tumor sample.

In comparison, both CLImAT and ASCAT provided robust purity estimate down to 20% simulated tumor purity, but substantially over-estimated tumor cellularity for the simulated low purity samples (**Figure 4C**). Both tools also failed to correctly resolve the correct ploidy of these low purity samples (**Figure 4A**). ASCAT provided results that were consistent with the ploidy estimates from sCNAphase for high purity samples, but at 40% and 20% purity, the estimates changed significantly for one and three of cell-lines respectively. CLImAT also showed the limitation in ploidy estimation at 40%, and it also misinterpreted the tumor ploidy of all the mixtures from HCC1954 and HCC2218. The sCNA predictions of CLImAT and ASCAT were self-consistent down to 40% purity for 3 of 4 samples, although with lower self-consistency scores than sCNAphase (**Figure 4D**). These results showcase the utility of the increased power offered by per-segment based haplotype counting strategy to resolve complex tumor genomes over the current state-of-the-art BAF based approach.

### The effect of sequencing and mapping artifacts

To test the tolerance of the methods to sequencing artifacts, we applied the tools to a matched tumor normal pair in which the tumor was an independent normal sample (i.e. a 0% tumor sample). In this case, the estimated tumor DNA purity should be 0%, and any sCNAs predicted are due to noise alone. For all four 0% tumour samples, sCNAphase identified < 0.1% tumor purity (**Figure** 4C; **Supplementary Table 3**). It did, however, identify spurious sCNAs present at this level of purity. On this basis, we can recommend that the sCNAphase segmentation should be disregarded if the estimated tumor cellularity is less than 1%. ASCAT and CLImAT reported tumor purity estimates of above 20% for these 0% tumor mixtures, which indicates that the copy number segmentation of these tools may be only reliable down to 40% purity (**Figure 4C**).

### Microarray analysis of cell-lines validates the sCNAphase results

The Cancer Cell Line Encyclopedia and the COSMIC Cell Line Project (34) provide an independent annotation of the mutations present in publically available cell-lines. The copy number profiles of the cell-lines analyzed by sCNAphase had been characterized as part of this COSMIC project, and were independently profiled using a PICNIC analysis of microarray data (35). This resource provides us with an independent annotation of the specific sCNAs in each of these cell-lines and allows us to investigate the capacity of sCNAphase to report on individual sCNAs.

To compare the annotations of these cell-lines (**Table 2**), the base ploidy for each cell-line were determined, by rounding up the ploidy estimates from SKY, flow cytometry and PICNIC to integers and taking the consensus values. In this process, hyper or hypo-teraploidy was round to tetraploidy; hypo-triploidy to triploidy. On this basis, HCC1187 was considered as triploid and all other cell-lines were treated as tetraploids. Any segment with a copy number greater than the ploidy was defined as an amplification, otherwise as a deletion. Sensitivity and specificity were calculated by counting the per base overlap for copy number gain or loss (Methods). This comparison revealed that the majority of events present in the COSMIC annotation of each cell line could be found across the range of mixtures for each cell-line using sCNAphase, when ploidy was properly assigned (**Figure** 5A, B). In the samples in which ploidy was incorrectly calculated (5% mixtures for HCC2218 and HCC1954), the capacity of sCNAphase to reflect the COSMIC results was greatly diminished. A similar comparison with the results from CLImAT showed that this approach failed to correctly profile any of the HCC2218 or HCC1954 mixtures (**Figure** 5A, B). ASCAT had comparable performance with sCNAphase for all four cell-lines in the samples that contained more than 40% tumor DNA, but the consistency with COSMIC segmentation significantly dropped at lower purity due to incorrect ploidy estimates. The comparison of the sCNAs identified by an array-based approach to those identified by sCNAphase, CLImAT and ASCAT, illustrate the ability of sCNAphase to identify valid copy number changes across a range of different simulated tumor purities.

**Figure 5.**
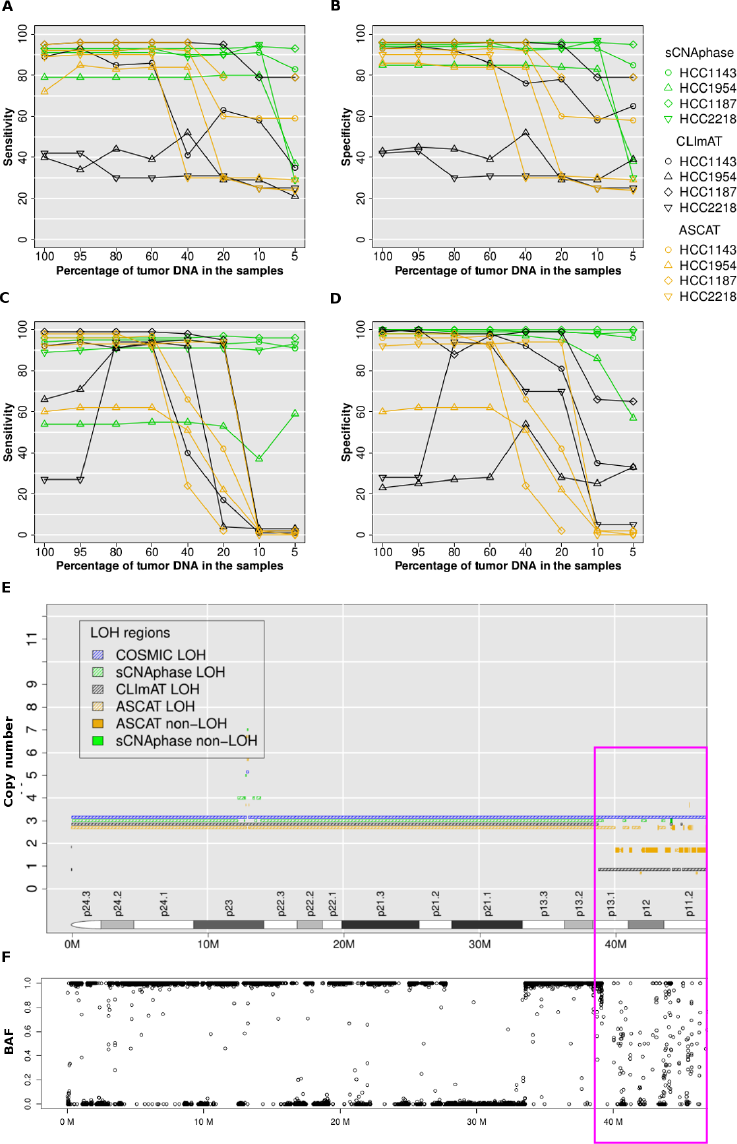
sCNAphase recapitulates the COSMIC annotation of these cell-lines across a range of purities. Consistency with COSMIC segmentation for copy number (A, B) and LOH (C,D) at varied tumor purity. For tumor samples at different tumor purity, the base-pair sensitivity in (A,C) and basepair specificity in (B,D), was calculated by overlapping sCNAphase, CLImAT and ASCAT segmentations estimated at a particular tumor purity with COSMIC segmentations based on 100% purity tumor samples. Each of the four cell-lines was represented a particular point shape as indicated in the legend. Results from sCNAphase, CLImAT and ASCAT are shown in green, black and orange respectively. The copy number segmentations of chr9 p-arm HCC1187 from COSMIC, sCNAphase, CLImAT and ASCAT were shown in (E) in blue, green, black and orange hash lines respectively. BAFs at this region in (F) support the loss of heterozygosity shown in COSMIC and CLImATsegmentation in p-arm except for the region highlighted in the pink box (39M-47M). For most of this 8M region, sCNAphase did not report copy number or LOH due to the highly variable BAFs which give merging errors (see Method). The complexity of this region can be also found from ASCAT predication at this region.

LOH events are a common feature of the cancer genome and have been previously linked to the inactivation of tumor suppressors (36, 37). Given the capacity of our approach to quantify and identify the haplotype of each chromosome, we assessed the ability of sCNAphase to identify the LOH events present in the COSMIC annotation of these cell lines (**Figure** 5C, D). This comparison showed sCNAphase identified approximately 90% of the regions of LOHs in COSMIC annotations of these cell-lines while producing few spurious results (except for HCC1954). Furthermore, sCNAphase was still able to identify the same regions of LOH in the low purity samples in which ploidy had been incorrectly assigned. CLImAT and ASCAT, however, showed very inconsistent LOH profile with COSMIC at low than 40% and 20% tumor purity.

One limitation for sCNAphase to reach even higher sensitivity was that sCNAphase provided no estimation at long regions with few SNPs or with highly variable depth profile (see Methods). For example, chr9p1-13 of HCC1187 was defined as a region of LOH by both COSMIC and CLImAT (**Figure** 5E); however sCNAphase only identified small islands of LOH (marked in pink) and left the majority as undetermined. When investigating the raw BAFs from this location (Figure 5F), the majority of the BAFs fell randomly between 0 to 1, producing a depth profile that was too confounding for sCNAphase to confidently resolve. The complexity of the copy number profile at this region was also recognized by ASCAT as revealed in the frequent switching between different copy number states.

In the analysis of HCC1954 there was a low degree of overlap between the regions of LOH identified by the three tools and those present in the COSMIC LOH annotation of the cell-line. Closer inspection indicated that these differences were due to large chromosomal regions predicted to be LOH in COSMIC, but which appear to be regions of high copy number and allelic imbalance, rather than regions of LOH (**Supplementary Figure** S6). We also found substantial differences between the copy number estimates from COSMIC and sCNAphase for multiple chromosomal arms for HCC1954 as well as differences between the ploidy estimates in COSMIC and those from sCNAphase and the previous SKY analyses (**Supplementary Figure** S4, S6). Given the inconsistent results, the COSMIC annotation of the HCC1954 may underestimate the ploidy of this cell-line (Table 2), and as a result, may have reported some regions of LOHs and focal deletions for HCC1954 that do not reflect the true copy number profile of this cell-line.

### Focal sCNAs identified by sCNAphase mirror those identified by microarray analysis

Given the clinical importance of recurrent focal amplifications in the cancer process, we assessed the performance of sCNAphase and CLImAT to detect the focal amplifications present in the COSMIC annotation for these four cell-lines, using the criteria of counting 50% reciprocal overlap for focal amplifications (Methods). A very stringent threshold was applied, which required focal amplifications to be in between 100 kb and 4 Mb as well as a copy number that was greater than twice the ploidy of the cell-line. It is worth noting that this is a more difficult copy number threshold for amplification than the one used in the previous section. In the analysis of the pure cell-lines, sCNAphase was able to detect the majority of the focal amplifications present in the COSMIC annotation of each of these cell-lines, with a sensitivity approximately twice that of CLImAT (**Table** 3; **Supplementary Table** S4).

For mixture samples, sCNAphase was still able to identify the majority of focal events at 5% tumor purity for HCC1187 and HCC1143, and 10% for HCC1954 and HCC2218. The drop in the 5% mixtures from HCC2218 and HCC1954 was due to the underestimation of the tumor ploidy in two samples. Despite this, sCNAphase was able to detect at least 12 copies of the pathologically relevant *ERBB2* in HCC1954 and HCC2218 across the entire cohort of mixtures (**Supplementary Table** S5), highlighting the diagnostic potential of sCNAphase (38, 39) at even 5% tumor DNA. Likewise, another focal peak of amplification, that was found to be recurrently altered in breast cancer (11q13) (40, 41), was detected in both the 100% HCC1143 and HCC1954 samples, as well as their corresponding 5% mixtures. The identification of focal events, including those that are well known to be clinically significant, across the full range of mixtures demonstrates the capacity of sCNAphase to identify pathologically relevant sCNAs in ultra-low purity samples.

We hypothesized that one reason for a low specificity (on average, only 41% and 31% of focal amplifications detected by sCNAphase and CLImAT respectively were validated by COSMIC) could be that COSMIC underestimates copy number of highly duplicated regions (due to fluorescence signal saturation) and so the 2 * ploidy threshold for declaring a focal amplification is not reached. To test this, we re-calculated specificity and sensitivity after increasing the sCNAphase detection threshold, but keeping the COSMIC threshold (**Supplementary Table S6**). As the sCNAphase threshold increased, the specificities almost doubled across all tumor purities, with a much smaller effect on sensitivity.

Homozygous deletions are another class of copy number mutation involved in the tumorigenic process (42-44). The COSMIC annotations show that there are 7 homozygous deletions larger than 100Kb in these four cell-lines. sCNAphase was able to consistently detect the 3 longer homozygous deletions from HCC1143 and HCC1187(**Supplementary Table S7**). The size of a focal deletion, more specifically, the number of germ-lines SNPs in the region limits sCNAphase from identifying shorter events. Our analysis of HCC1954 did not detect the longer deletion from chr22 of HCC1954; however analysis of the raw sequencing data suggest that this deletion may be shorter than the 332kb listed in the COSMIC annotation, as the read depth at the flanking region is significantly higher than the deleted region **(Supplementary Figure S7**). Despite this, sCNAphase did not identify any false positives and was able to consistently identify these deletions at minimal levels of tumor purity. In contrast, CLImAT failed to identify any of homozygous deletions present in COSMIC, and it classified a few other regions as homozygous deletions that were not present in either the COSMIC or sCNAphase results.

### sCNAphase identifies Copy Neutral / Amplified Regions of LOH in low purity samples

Examination of the pure cell-lines with sCNAphase, revealed a substantial fraction of the tumor genome that had undergone LOH and contained two or more copies (**Table 2**). This is a compound event which requires at least a deletion of one haplotype as well as amplification of the other haplotype. Analysis of these regions revealed a number of loci that were present in multiple unrelated cell-lines, including chr5q and chr17p as well as parts of chr17q, all of which had been previously found to undergo recurrent CN-LOH in cancer (37, 45-47) (**Supplementary Figure S4**). One of these regions contains the tumor suppressor *TP53*, the gene most commonly altered by CN-LOH (45, 48). We observed homozygous somatic mutations in *TP53* for three of the cell-lines (**Supplementary Table S8**), which are likely to be disruptive. In the remaining cell-line, two heterozygous germline SNPs were identified (**Supplementary Table S9**), one of which, R283C has been previously shown to increase a carrier’s risk of developing cancer (45,49,50). It is likely that the loss of the wild type allele and the amplification of the deleterious SNP increase this risk. These results demonstrate the power offered by HTS and sCNAphase to characterize the biological impacts of a complex mutation in low purity samples.

## DISCUSSION

Although somatic copy number alterations are a well-established driver of cancer, the capacity to identify these mutations are impacted by a number of issues including varying levels of tumor purity and frequent changes in tumor ploidy. As a result, the majority of methods designed to characterize the sCNAs in the cancer genome are unable to accurately profile a significant fraction of primary tumor samples. In this study, we have shown that by taking a haplotype-based approach, sCNAphase can overcome these issues and reliably characterize both genome-wide, chromosome-arm-wide as well as focal copy number changes present in a tumor genome across a range of tumor purities.

Comparison of the results from the sCNAphase to those from SKY and flow cytometry, illustrated the ability of sCNAphase to correctly reflect the genome-wide changes in ploidy. Using a range of mixtures, to simulate the challenges posed by low-cellularity primary tumors, we were able to recapitulate the changes in ploidy seen in the analysis of the pure cell-lines, across the spectrum of tumor purities. Comparison of copy number and LOH segmentations obtained from a high density microarray at 100% purity to those reported by sCNAphase across a range of mixture samples demonstrated the high specificity and sensitivity of the method down to 10% tumor purity.

Equally importantly, samples without any tumor DNA were predicted to have < 0.1% tumor cellularity by sCNAphase, whereas both of the other approaches tested reported at least 20% tumor DNA. sCNAphase has this in-built robustness to sequencing and mapping artifacts because it models the observed regional tumor depth data (including total depth and haplotype-specific depth), conditional on normal depth data from the same region.

To focus on characterizing the copy number profile of low purity tumors, we have made the simplifying assumption that there is a dominant tumor clone with low heterogeneity in the tumor biopsy. Regions with heterogeneous copy number between clones with similar abundance would lead to a failure of the merging test statistic, and thus these regions would likely be excluded.

The accurate characterization of the copy number profile in low cellularity samples as well as the identification of mutations in cancer genes, is suggestive of the potential clinical utility of this tool. In addition to low purity tumors, a potential application of our haplotype-based methodology would be in studies that aim to profile the copy number changes in a tumor through the analysis of circulating tumor DNA. Circulating tumor DNA, has been previously shown to contain mutations present in the tumor genome, but tumor DNA in circulation is mixed with DNA from normal cells and the tumor purity is frequently low (51). While sCNAphase can successfully profile low purity mixtures, it is likely that in order to realize this, an optimized version of the tool will need to be developed.

## ACKNOWLEDGEMENT

The work was supported by UQ-CSC scholarship awarded to Wenhan Chen. We thank Haojing Shao for analyzing the patterns of the somatic copy number alterations, Dr Sarah Song for collecting the sequencing data and Dr Alison Anderson for reviewing on the current study, Dr Paul Edwards for granting us the permission of editing the SKY data, and Dr. Peter Van Loo and Dr. Kerstin Haase for the advices on how to run ASCAT for HTS data. We appreciate the WGS data from TCGA consortium, in particular, Dr Adam Ewing for creating the tumor-normal mixtures for HCC1143 and HCC1954. We would also like to thank Illumina BaseSpace for providing the WGS data for HCC1187 and HCC2218, COSMIC server for providing the array-based copy number profiles.

## FUNDING

The work was supported by Cancer Council Queensland grant [APP1085786] awarded to Dr. Lachlan J.M. Coin and UQ-CSC scholarship awarded to Wenhan Chen. Funding for open access charge: Cancer Council Queensland.

## REFERENCES

1. Beroukhim, R., Mermel, C.H., Porter, D., Wei, G., Raychaudhuri, S., Donovan, J., Barretina, J., Boehm, J.S., Dobson, J., Urashima, M. et al. (2010) The landscape of somatic copy-number alteration across human cancers. Nature, 463, 899-905.

2. Bambury, R.M., Bhatt, A.S., Riester, M., Pedamallu, C.S., Duke, F., Bellmunt, J., Stack, E.C., Werner, L., Park, R., Iyer, G. et al. (2015) DNA copy number analysis of metastatic urothelial carcinoma with comparison to primary tumors. BMC Cancer, 15, 242-242.

3. Lee, A.J.X., Endesfelder, D., Rowan, A.J., Walther, A., Birkbak, N.J., Futreal, P.A., Downward, J., Szallasi, Z., Tomlinson, I.P.M., Howell, M. et al. (2011) Chromosomal instability confers intrinsic multidrug resistance. Cancer research, 71, 1858-1870.

4. Carter, S.L., Eklund, A.C., Kohane, I.S., Harris, L.N. and Szallasi, Z. (2006) A signature of chromosomal instability inferred from gene expression profiles predicts clinical outcome in multiple human cancers. Nature genetics, 38, 1043-1048.

5. Hieronymus, H., Schultz, N., Gopalan, A., Carver, B.S., Chang, M.T., Xiao, Y., Heguy, A., Huberman, K., Bernstein, M., Assel, M. et al. (2014) Copy number alteration burden predicts prostate cancer relapse. Proceedings of the National Academy of Sciences, 111, 11139-11144.

6. Sheffer, M., Bacolod, M.D., Zuk, O., Giardina, S.F., Pincas, H., Barany, F., Paty, P.B., Gerald, W.L., Notterman, D.a. and Domany, E. (2009) Association of survival and disease progression with chromosomal instability: a genomic exploration of colorectal cancer. Proceedings of the National Academy of Sciences, 106, 7131-7136.

7. Zack, T.I., Schumacher, S.E., Carter, S.L., Cherniack, A.D., Saksena, G., Tabak, B., Lawrence, M.S., Zhang, C.-Z., Wala, J., Mermel, C.H. et al. (2013) Pan-cancer patterns of somatic copy number alteration. Nat Genet, 45, 1134-1140.

8. Cho, Y.-J., Tsherniak, A., Tamayo, P., Santagata, S., Ligon, A., Greulich, H., Berhoukim, R., Amani, V., Goumnerova, L., Eberhart, C.G. et al. (2011) Integrative genomic analysis of medulloblastoma identifies a molecular subgroup that drives poor clinical outcome. Journal of the American Society of Clinical Oncology, 29, 1424-1430.

9. Chin, K., DeVries, S., Fridlyand, J., Spellman, P.T., Roydasgupta, R., Kuo, W.-L., Lapuk, A., Neve, R.M., Qian, Z., Ryder, T. et al. (2006) Genomic and transcriptional aberrations linked to breast cancer pathophysiologies. Cancer cell, 10, 529-541.

10. Curtis, C., Shah, Sp. Chin, S.-F., Turashvili, G., Rueda, O.M., Dunning, M.J., Speed, D., Lynch, A.G., Samarajiwa, S., Yuan, Y. et al. (2012) The genomic and transcriptomic architecture of 2,000 breast tumours reveals novel subgroups. Nature, 486, 346-352.

11. Slamon, D.J., Leyland-Jones, B., Shak, S., Fuchs, H., Paton, V., Bajamonde, A., Fleming, T., Eiermann, W., Wolter, J., Pegram, M. et al. (2001) Use of Chemotherapy plus a Monoclonal Antibody against HER2 for Metastatic Breast Cancer That Overexpresses HER2. New England Journal of Medicine, 344, 783-792.

12. Bast, R.C., Ravdin, P., Hayes, D.F., Bates, S., Fritsche, H., Jessup, J.M., Kemeny, N., Locker, G.Y., Mennel, R.G. and Somerfield, M.R. (2001) 2000 Update of Recommendations for the Use of Tumor Markers in Breast and Colorectal Cancer: Clinical Practice Guidelines of the American Society of Clinical Oncology*. Journal of Clinical Oncology, 19, 1865-1878.

13. Carter, S.L., Cibulskis, K., Helman, E., McKenna, A., Shen, H., Zack, T., Laird, P.W., Onofrio, R.C., Winckler, W., Weir, B.a. et al. (2012) Absolute quantification of somatic DNA alterations in human cancer. Nature biotechnology, 30, 413-421.

14. Aran, D., Sirota, M. and Butte, A.J. (2015) Systematic pan-cancer analysis of tumour purity. Nat Commun, 6.

15. Sirivatanauksorn, V., Sirivatanauksorn, Y., Gorman, P.A., Davidson, J.M., Sheer, D., Moore, P.S., Scarpa, A., Edwards, P.A.W. and Lemoine, N.R. (2001) Non-random chromosomal rearrangements in pancreatic cancer cell lines identified by spectral karyotyping. International journal of cancer, 91, 350-358.

16. Storlazzi, C.T., Lonoce, A., Guastadisegni, M.C., Trombetta, D., D’Addabbo, P., Daniele, G., L’Abbate, A., Macchia, G., Surace, C., Kok, K. et al. (2010) Gene amplification as double minutes or homogeneously staining regions in solid tumors: origin and structure. Genome research, 20, 1198-1206.

17. Stephens, P.J., Greenman, C.D., Fu, B., Yang, F., Bignell, G.R., Mudie, L.J., Pleasance, E.D., Lau, K.W., Beare, D., Stebbings, L.a. et al. (2011) Massive genomic rearrangement acquired in a single catastrophic event during cancer development. Cell, 144, 27-40.

18. Schröck, E., Manoir, S.d., Veldman, T., Schoell, B., Wienberg, J., Ferguson-Smith, M.A., Ning, Y., Ledbetter, D.H., Bar-Am, I., Soenksen, D. et al. (1996) Multicolor Spectral Karyotyping of Human Chromosomes. Science, 273, 494-497.

19. Zhao, X., Li, C., Paez, J.G., Chin, K., Jänne, P.A., Chen, T.-H., Girard, L., Minna, J., Christiani, D. and Leo, C. (2004) An integrated view of copy number and allelic alterations in the cancer genome using single nucleotide polymorphism arrays. Cancer research, 64, 3060-3071.

20. Conrad, D.F., Pinto, D., Redon, R., Feuk, L., Gokcumen, O., Zhang, Y., Aerts, J., Andrews, T.D., Barnes, C., Campbell, P. et al. (2010) Origins and functional impact of copy number variation in the human genome. Nature, 464, 704-712.

21. Krijgsman, O., Carvalho, B., Meijer, G.a., Steenbergen, R.D.M. and Ylstra, B. (2014) Focal chromosomal copy number aberrations in cancer-Needles in a genome haystack. Biochimica et biophysica acta, 1843, 2698-2704.

22. Pinto, D., Darvishi, K., Shi, X., Rajan, D., Rigler, D., Fitzgerald, T., Lionel, A.C., Thiruvahindrapuram, B., MacDonald, J.R. and Mills, R. (2011) Comprehensive assessment of array-based platforms and calling algorithms for detection of copy number variants. Nature biotechnology, 29, 512-520.

23. Van Loo, P., Nordgard, S.H., Lingjærde, O.C., Russnes, H.G., Rye, I.H., Sun, W., Weigman, V.J., Marynen, P., Zetterberg, A. and Naume, B. (2010) Allele-specific copy number analysis of tumors. Proceedings of the National Academy of Sciences, 107, 16910-16915.

24. Goya, R., Sun, M.G., Morin, R.D., Leung, G., Ha, G., Wiegand, K.C., Senz, J., Crisan, A., Marra, M.A. and Hirst, M. (2010) SNVMix: predicting single nucleotide variants from next-generation sequencing of tumors. Bioinformatics, 26, 730-736.

25. Boeva, V., Popova, T., Bleakley, K., Chiche, P., Cappo, J., Schleiermacher, G., Janoueix-Lerosey, I., Delattre, O. and Barillot, E. (2012) Control-FREEC: a tool for assessing copy number and allelic content using next-generation sequencing data. Bioinformatics, 28, 423-425.

26. Mayrhofer, M., DiLorenzo, S. and Isaksson, A. (2013) Patchwork: allele-specific copy number analysis of whole-genome sequenced tumor tissue. Genome Biol, 14, R24.

27. Altshuler, D., Durbin, R., Abecasis, G., Bentley, D., Chakravarti, A., Clark, A., Donnelly, P., Eichler, E., Flicek, P. and Gabriel, S. (2012) An integrated map of genetic variation from 1,092 human genomes. NATURE, 491, 56-65.

28. Delaneau, O., Howie, B., Cox, A.J., Zagury, J.-F. and Marchini, J. (2013) Haplotype estimation using sequencing reads. American journal of human genetics, 93, 687-696.

29. Wilks, C., Cline, M.S., Weiler, E., Diehkans, M., Craft, B., Martin, C., Murphy, D., Pierce, H., Black, J., Nelson, D. et al. (2014) The Cancer Genomics Hub (CGHub): overcoming cancer through the power of torrential data. Database : the journal of biological databases and curation, 2014.

30. Li, H., Handsaker, B., Wysoker, A., Fennell, T., Ruan, J., Homer, N., Marth, G., Abecasis, G., Durbin, R. and Subgroup, G.P.D.P. (2009) The Sequence Alignment/Map format and SAMtools. Bioinformatics, 25, 2078-2079.

31. Rabiner, L.R. (1989) A tutorial on hidden Markov models and selected applications in speech recognition. Proceedings of the IEEE, 77, 257-286.

32. Forney, G.D. (1973) The viterbi algorithm. Proceedings of the IEEE, 61, 268-278.

33. Coin, L.J., Asher, J.E., Walters, R.G., Moustafa, J.S.E.-S., de Smith, A.J., Sladek, R., Balding, D.J., Froguel, P. and Blakemore, A.I. (2010) cnvHap: an integrative population and haplotype–based multiplatform model of SNPs and CNVs. Nature methods, 7, 541-546.

34. Forbes, S.A., Beare, D., Gunasekaran, P., Leung, K., Bindal, N., Boutselakis, H., Ding, M., Bamford, S., Cole, C., Ward, S. et al. (2015) COSMIC: exploring the world’s knowledge of somatic mutations in human cancer. Nucleic Acids Research, 43, D805-D811.

35. Greenman, C.D., Bignell, G., Butler, A., Edkins, S., Hinton, J., Beare, D., Swamy, S., Santarius, T., Chen, L. and Widaa, S. (2010) PICNIC: an algorithm to predict absolute allelic copy number variation with microarray cancer data. Biostatistics, 11, 164-175.

36. Side, L., Taylor, B., Cayouette, M., Conner, E., Thompson, P., Luce, M. and Shannon, K. (1997) Homozygous Inactivation of the NF1 Gene in Bone Marrow Cells from Children with Neurofibromatosis Type 1 and Malignant Myeloid Disorders. New England Journal of Medicine, 336, 1713-1720.

37. Tuna, M., Knuutila, S. and Mills, G.B. (2009) Uniparental disomy in cancer. Trends in Molecular Medicine, 15, 120-128.

38. Kao, J., Salari, K., Bocanegra, M., Choi, Y.-L., Girard, L., Gandhi, J., Kwei, K.A., Hernandez-Boussard, T., Wang, P., Gazdar, A.F. et al. (2009) Molecular Profiling of Breast Cancer Cell Lines Defines Relevant Tumor Models and Provides a Resource for Cancer Gene Discovery. PLoS ONE, 4, e6146.

39. Neve, R.M., Chin, K., Fridlyand, J., Yeh, J., Baehner, F.L., Fevr, T., Clark, L., Bayani, N., Coppe, J.-P., Tong, F. et al. (2006) A collection of breast cancer cell lines for the study of functionally distinct cancer subtypes. Cancer Cell, 10, 515-527.

40. Desnoyers, L.R., Pai, R., Ferrando, R.E., Hotzel, K., Le, T., Ross, J., Carano, R., D’Souza, A., Qing, J., Mohtashemi, I. et al. (2007) Targeting FGF19 inhibits tumor growth in colon cancer xenograft and FGF19 transgenic hepatocellular carcinoma models. Oncogene, 27, 85-97.

41. Sawey, Eric T., Chanrion, M., Cai, C., Wu, G., Zhang, J., Zender, L., Zhao, A., Busuttil, Ronald W., Yee, H., Stein, L. et al. (2011) Identification of a Therapeutic Strategy Targeting Amplified FGF19 in Liver Cancer by Oncogenomic Screening. Cancer Cell, 19, 347-358.

42. Li, J., Yen, C., Liaw, D., Podsypanina, K., Bose, S., Wang, S.I., Puc, J., Miliaresis, C., Rodgers, L., McCombie, R. et al. (1997) PTEN, a Putative Protein Tyrosine Phosphatase Gene Mutated in Human Brain, Breast, and Prostate Cancer. Science, 275, 1943-1947.

43. Cairns, P., Polascik, T.J., Eby, Y., Tokino, K., Califano, J., Merlo, A., Mao, L., Herath, J., Jenkins, R., Westra, W. et al. (1995) Frequency of homozygous deletion at p16/CDKN2 in primary human tumours. Nat Genet, 11, 210-212.

44. Knudson, A.G., Hethcote, H.W. and Brown, B.W. (1975) Mutation and childhood cancer: a probabilistic model for the incidence of retinoblastoma. Proceedings of the National Academy of Sciences, 72, 5116-5120.

45. Murthy, S.K., DiFrancesco, L.M., Ogilvie, R.T. and Demetrick, D.J. (2002) Loss of Heterozygosity Associated with Uniparental Disomy in Breast Carcinoma. Mod Pathol, 15, 1241-1250.

46. Pedersen, B.S., Konstantinopoulos, P.A., Spillman, M.A. and De, S. (2013) Copy neutral loss of heterozygosity is more frequent in older ovarian cancer patients. Genes, Chromosomes and Cancer, 52, 794-801.

47. Lips, E.H., de Graaf, E.J., Tollenaar, R., van Eijk, R., Oosting, J., Szuhai, K., Karsten, T., Nanya, Y., Ogawa, S., van de Velde, C.J. et al. (2007) Single nucleotide polymorphism array analysis of chromosomal instability patterns discriminates rectal adenomas from carcinomas. The Journal of Pathology, 212, 269-277.

48. Mullighan, C.G., Goorha, S., Radtke, I., Miller, C.B., Coustan-Smith, E., Dalton, J.D., Girtman, K., Mathew, S., Ma, J., Pounds, S.B. et al. (2007) Genome-wide analysis of genetic alterations in acute lymphoblastic leukaemia. Nature, 446, 758-764.

49. Keller, G., Vogelsang, H., Becker, I., Plaschke, S., Ott, K., Suriano, G., Mateus, A.R., Seruca, R., Biedermann, K. and Huntsman, D. (2004) Germline mutations of the E-cadherin (CDH1) and TP53 genes, rather than of RUNX3 and HPP1, contribute to genetic predisposition in German gastric cancer patients. Journal of medical genetics, 41, e89-e89.

50. Manoukian, S., Peissel, B., Pensotti, V., Barile, M., Cortesi, L., Stacchiotti, S., Terenziani, M., Barbera, F., Pasquini, G., Frigerio, S. et al. (2007) Germline mutations of TP53 and BRCA2 genes in breast cancer/sarcoma families. European Journal of Cancer, 43, 601-606.

51. Newman, A.M., Bratman, S.V., To, J., Wynne, J.F., Eclov, N.C.W., Modlin, L.A., Liu, C.L., Neal, J.W., Wakelee, H.A. and Merritt, R.E. (2014) An ultrasensitive method for quantitating circulating tumor DNA with broad patient coverage. Nature medicine.

52. Grigorova, M., Lyman, R.C., Caldas, C. and Edwards, P.A.W. (2005) Chromosome abnormalities in 10 lung cancer cell lines of the NCI-H series analyzed with spectral karyotyping. Cancer Genetics and Cytogenetics, 162, 1-9.

